# Evaluation of the tyrosine and dopamine serum level in experimental infected BALB/c mice with chronic toxoplasmosis

**DOI:** 10.1101/2020.12.17.423217

**Authors:** Mehdi Mirzaeipour, Fattaneh Mikaeili, Qasem Asgari, Mohammad Nohtani, Sajad Rashidi, Mohammad Saleh Bahreini

## Abstract

**Background:** *Toxoplasma* parasite alters the transduction of neurotransmitter signals and leads to changes in the level of brain neurotransmitters including tyrosine and dopamine, so, behavior changes can occur in infected hosts. Based on this concept, this study was conducted for evaluation of the tyrosine and dopamine serum level in infected mice with chronic toxoplasmosis.

**Materials and methods:** *Toxoplasma gondii* (Prugniaud strain II) was injected intra-peritoneal into BALB/c mice to induce chronic toxoplasmosis. Modified agglutination test (MAT), polymerase chain reaction (PCR), and microscopic methods were conducted to confirm the induction of chronic toxoplasmosis. The infected mice sera were separated at days 40, 50, 60, 70, and 80 for evaluation of tyrosine and dopamine serum level using High-performance Liquid Chromatography (HPLC).

**Results:** Microscopic methods confirmed the formation of the *Toxoplasma* cysts in mice tissues. Inducing chronic toxoplasmosis is also confirmed by MAT, PCR and histological methods. HPLC results indicated a decrease in serum tyrosine level at days 40 in infected mice in comparison to control and the levels were too low to be measured at other times. However, a significantly high serum dopamine level was observed that gradually increased after parasite inoculation.

**Conclusions:** No detection of tyrosine level in most of the sample groups is probably related to the very low concentration of tyrosine in sera. However, low concentration of tyrosine at days 40 and increase of dopamine in most of the sample groups suggests the production of dopamine from tyrosine due to the presence of *Toxoplasma* in infected mice.

## Introduction

Toxoplasmosis is an infectious disease that caused by *Toxoplasma gondii;* an intracellular protozoan belonging to the Phylum Apicomplexa. Toxoplasmosis is highly prevalent throughout the world in human and warm-blooded animals. In congenital type, as an important type of toxoplasmosis, the transmission of the parasite from the placenta can lead to inducing severe complications in the fetus. Ophthalmological and neurological disorders are considered as remarkable outcomes of the congenital form (1, 2).

Acquired toxoplasmosis is classified into symptomatic and asymptomatic types. The symptomatic type usually occurs in individuals with HIV and other immune system disorders(3–5). Toxoplasmic encephalitis is considered a predictable outcome of neurological disorders in HIV positive individuals (6). Several recent studies have revealed the association of toxoplasmosis and neurodegenerative disorders including Parkinson, Multiple Sclerosis and Alzheimer (7–10).

Particular researches pointed out the relationship of host’s behavior changes and different microorganisms including viruses and parasites such as *T. gondii* (11–13). Although the main reason for these behavioral changes has not been precisely indicated so far, the formation of *Toxoplasma* cysts in the central nervous system (CNS) can be related to these changes. Cyst stage as a dormant form of *Toxoplasma* parasite is composed in muscle tissues and central nervous system (CNS). The release of bradyzoites due to spontaneous tissue cysts rupture leads to the continual high titer of the host serum antibodies (14, 15).

A few investigations declared that *Toxoplasma* parasite alters the transduction of neurotransmitter signals. They believe that the parasite overexpresses the encoding genes of tyrosine hydroxylase. Tyrosine hydroxylase is considered as a rate-limiting enzyme for producing dopamine (16). Also, the results of another study showed use of dopamine antagonists in psychiatric patients infected with toxoplasmosis decreases the behavioral changes in these patients (17).

Based on mentioned above, this study was conducted to measure tyrosine and dopamine serum level in experimentally infected mice with chronic toxoplasmosis using HPLC.

## Materials and methods

### Ethics approval

All experiments with mice were performed by following the guidelines of the Institutional Animal Care and Committee on Ethics of Animal Experimentation and approved by the Ethics Committee of Shiraz University of Medical Sciences (IR.SUMS.REC.1397.S653).

### Mice

In this study, seventy 6–8-week female inbred BALB/c mice (weight 30-35g) were provided by Pasteur Institute, Tehran, Iran. All used animals in this study were housed in temperature-controlled accommodation (22 ± 2 °C, 40-60% humidity), provided with water and fed with standard rodent dried food. The animals were kept at the Laboratory Animal Center of Shiraz University of Medical Sciences, Shiraz, Iran.

### Parasite preparation

*T. gondii* parasite (Prugniaud strain II (PRU- II)) was obtained from the brains of infected BALB/c mice that provided by the Department of Parasitology and Mycology, Mazandaran University of Medical Sciences. Briefly, 4 infected mice were sacrificed using phenol anesthesia. The brains of the infected animals were aseptically removed in sterile conditions. The brains were homogenized in a homogenizer solution containing 1% penicillin/streptomycin (Gibco Company). The homogenate was centrifuged at 100× g for 3 min at 4 °C to decrease the large debris. Then, the supernatant was centrifuged at 2000× g for 6 min at 4 °C. After re-suspension and dilution of the obtained pellet in normal saline, the cysts were counted.

### Inducing experimental chronic toxoplasmosis in mice (pilot group)

As a pilot study, five female mice were selected for induction of experimental chronic toxoplasmosis. For this aim, each mouse was infected intra-peritoneal using 200 μL of normal saline containing 10 cysts. After 2 months of parasite injection, the mice were sacrificed. Experimental chronic toxoplasmosis and cyst formation were confirmed using techniques including modified agglutination test (MAT) on the sera of mice, presence of parasites in brain and eye using direct and pathological smears polymerase chain reaction (PCR).

### MAT

MAT is considered as a sensitive and specific serologic test to measure antibodies against *T. gondii* in different hosts (18). The extracted *T. gondii* tachyzoites (RH strain) from the peritoneum of infected mice were used as antigen in MAT. Extracted tachyzoites were kept and fixed in 2% formalin at 4 °C overnight. In continue, formalin was washed using PBS and then, resulting pellet was re-suspended in alkaline borate buffer (pH 8.7) containing 0.4% Bovine Serum Albumin (BSA), 0.2% Sodium azide and Evans blue dye (40 μg/ml) to a final concentration including 2×108/ml parasites and stored at 4 °C.

The sera dilutions (1:20 to 1:2560) were provided with alkaline borate buffer in wells of U-bottom-shape 96 well microplate. Positive and negative sera were used and diluted similar to what was done for samples. After, 20 μl of 2-Mercaptoethanol (1/1000 V/V) was added to each wells and the plate was quietly shacked for 10 minutes.

40 μL of prepared antigen containing tachyzoites from the previous step was added to each well. The plate was incubated at room temperature for 24 h. and the results were read. In MAT, production of a blue carpet shape at the bottom of the wells including serum titers more than 1:20 are considered as positive results.

### Preparation of direct and pathology smears

Mice in the pilot study were sacrificed using phenol anesthesia. Mice brains and eyes were aseptically gathered and fixed in 10% buffered formalin. The resultant pieces are dehydrated with graded alcohols and then embedded in paraffin blocks. Sections of 5μm in thickness are prepared on slides and stained by hematoxylin and eosin and observed by light microscopy (19). Also impresser smears were prepared from the brain samples stained by Giemsa staining.

### PCR

DNA extraction from the brains was performed using a commercial kit (QIAGEN 28106, US). Specific primers were designed based on a genomic fragment of the B1 gene, with approximately 35 repeated sequences in the *Toxoplasma* genomes. Used primers (Forward: 5′-GGA ACT GCA TCC GTT CAT GAG-3′ and Reverse: 5′-TCT TTA AAG CGT TCG TGG TC-3) with high accuracy were able to detect a tachyzoite in samples (20).

The amplification program was set to start with pre-denaturation at 94 °C for 5 min, then 40 cycles of denaturation at 94 °C for 30 s, annealing at 57 °C for 45 s, and extension at 72 °C for 1 min, followed by a final extension stage at 72 °C for 10 min (20).

In PCR products electrophoresis stage, 5 μL of loading buffer was added to 10 μL of the final PCR product and was run to electrophoresis on 1.5% agarose gel and visualized under UV light with ethidium bromide.

### Inducing experimental chronic toxoplasmosis in mice (sample groups)

Sixty female BALB/c mice with the same age and weight as the pilot group were selected and divided into two groups (50 mice as samples and 10 mice as control groups). All mice in sample groups were infected the same as the pilot group. Also, 200 μL of normal saline as a placebo was injected intra-peritoneal into each mouse in the control group. In this study, BALB/c mice were maintained in the Institute of Comparative-Experimental Medicine of Shiraz University of Medical Sciences.

In the next step, the blood samples were taken from mice hearts at days 40, 50, 60, 70 and 80 after inducing chronic toxoplasmosis. The sera were separated and stored at −70 °C to measure tyrosine and dopamine serum level using HPLC system.

### Evaluation of the tyrosine and dopamine serum levels using HPLC

#### The setting of the HPLC system

Knauer HPLC system (Germany) in this study contained an automatic sampler, quaternary pump, degasser, and a detector. EZchrome software (Knauer/Germany) was used for analyzing the obtained signals. The used column was silica based, C18 (150×3.9 mm) that are based on 4 μm particle technology. The mobile phase was determined 0.05% formic acid and acetonitrile (90:10, v/v) with a 1 ml/min flow rate. All samples were injected into the HPLC with equal volumes (50 μl) at 225 nm and 236 nm wavelength for tyrosine and dopamine respectively. Retention time for tyrosine determined 3.6min & dopamine 3.4min.

#### Preparation of standard solutions

For preparation of standard solutions and setting the HPLC, 0.01 gr of tyrosine and dopamine were mixed separately with 0.5% HClO_4_ (v/v) for achievement of different concentrations including 1, 0.5, 0.125, 0.0625, and 0.03125 μg/ml. Each of the standard solutions was injected 3 times into the HPLC system and then all related curves were obtained.

#### Preparation of sample solutions

50 μl of each serum was mixed with 0.5% HClO4 (v/v) with equal volumes. Then, 50 μl of the obtained solution was injected into the HPLC. Related curves of all samples were obtained within 12 minutes.

#### Statistical analysis

SPSS statistics software (ver. 24, Chicago, IL, USA) and Mann-Whitney nonparametric test were used to analyze the obtained data. P-value ≤ 0.05 was statistically significant.

## Results

### MAT

MAT results showed that all mice sera had positive titers more than 1:20 in the pilot group. Since the ratio of 1:20 is considered as a positive cut off in interpretation of MAT results, induction of chronic toxoplasmosis was confirmed in the pilot group.

### Preparation of direct and pathology smears

Pathology smears results showed that after 50 days, *Toxoplasma* cysts were composed in mice brain and eye in the pilot group. Also, cysts were seen in direct smears with Giemsa staining. Our investigations also showed that the cysts are composed in mice brain and eye in sample groups (Figure 1).

**Figure 1.**
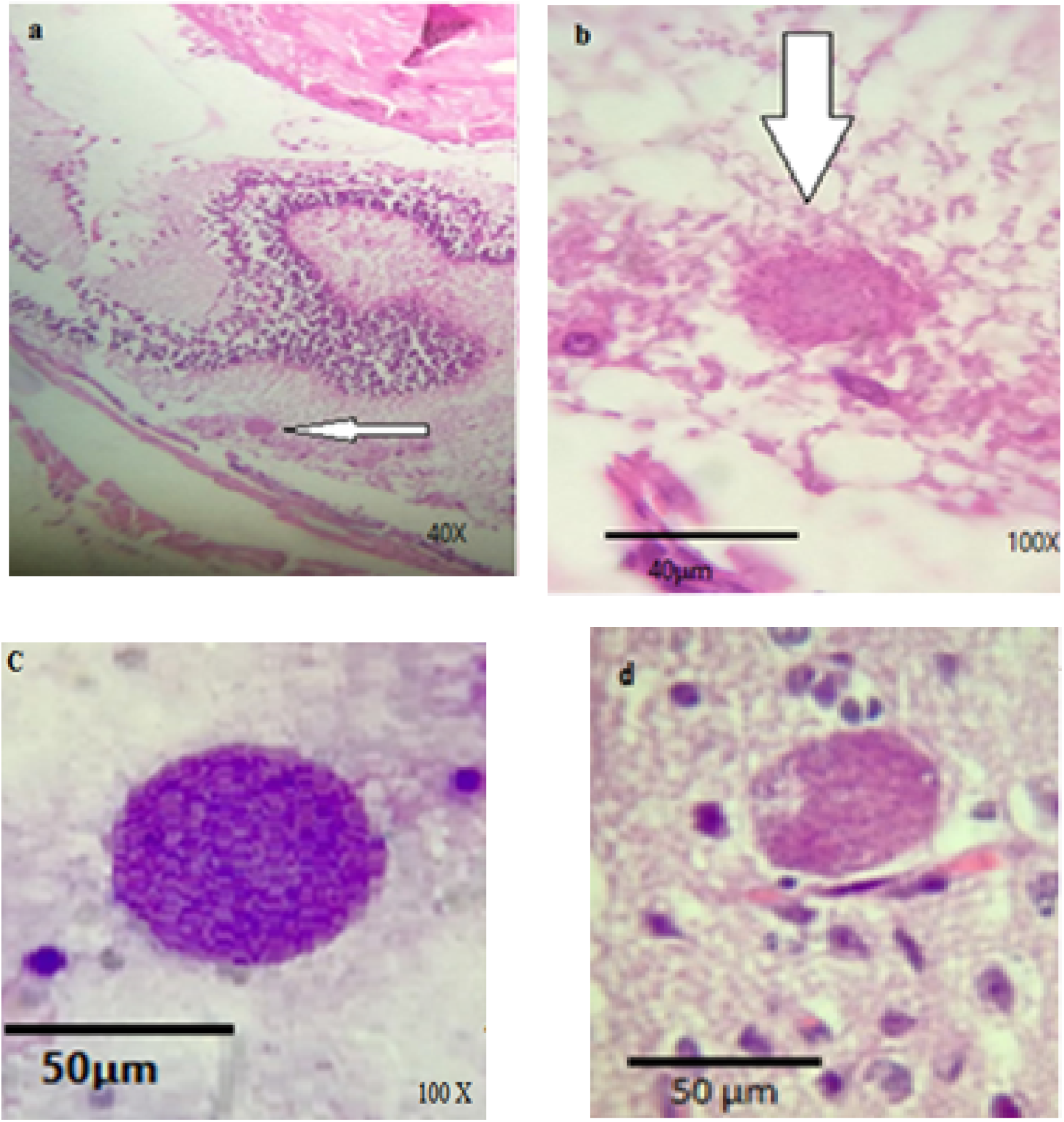
*Toxoplasma* cysts in sample groups: a, b) Eye tissue at days 50 with H&E staining, c) Direct smear of brain with Giemsa staining at days 50 d) Brain tissue pilot group with H&E staining

### PCR

PCR was developed to confirm the induction of chronic toxoplasmosis in the pilot group, the presence of *T. gondii* cysts in the mice brain, and microscopic results. By using the B1-F and B1-R primers, the positive samples for *T. gondii* presented 195-200 bp bands (Figure 2).

**Figure 2.**
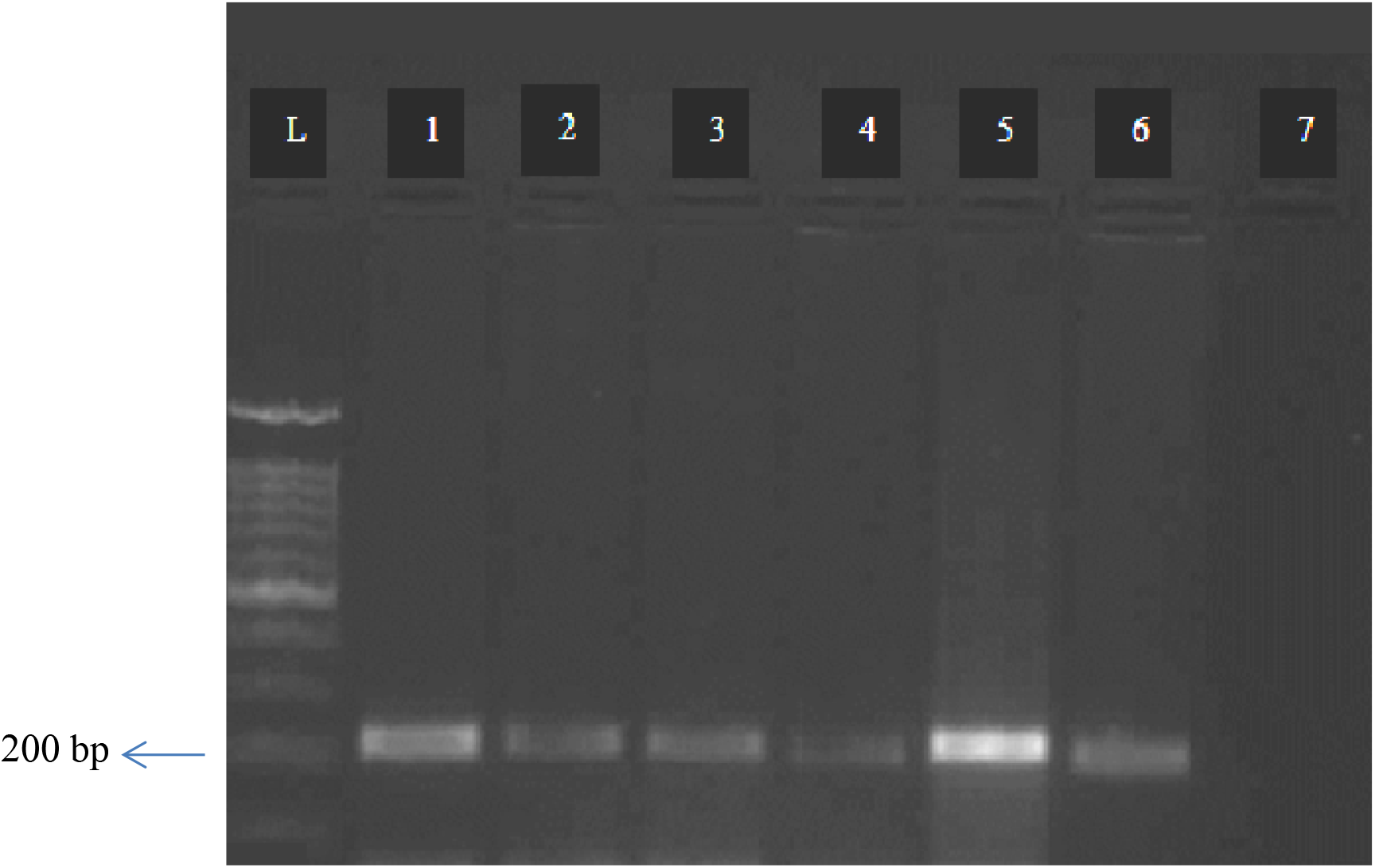
PCR results in mice brain (pilot groups): L) ladder 1000 bp, 1, 5) positive controls (*Toxoplasma* tachyzoites), 2, 3, 4, 6) samples, 7) negative control

### Evaluation of the tyrosine and dopamine serum level using HPLC

International Council for Harmonization guideline (ICH) was used for evaluation and validation of analytical procedures in the HPLC system. Resolution and injection precision, tailing factor, and the number of theoretical plates were checked as important factors for the performance of the system. For checking the non-interference between the level of tyrosine and dopamine in sera and other products such as mobile phase, placebo solution (a mixture of all the ingredients except sera), diluents, the standard solutions, and the sample solutions were injected separately into the HPLC system.

After preparation of tyrosine and dopamine standards in different concentrations and injection into the HPLC, standard curves were achieved. After calculation of the present area under obtained standard curves and drawing the calibration curves, tyrosine and dopamine standard equations were obtained. Finally, the concentration of tyrosine and dopamine in samples was measured based on the calculation of the present area under obtained sample curves and using the standard equations (Figures 3, 4).

**Figure 3.**
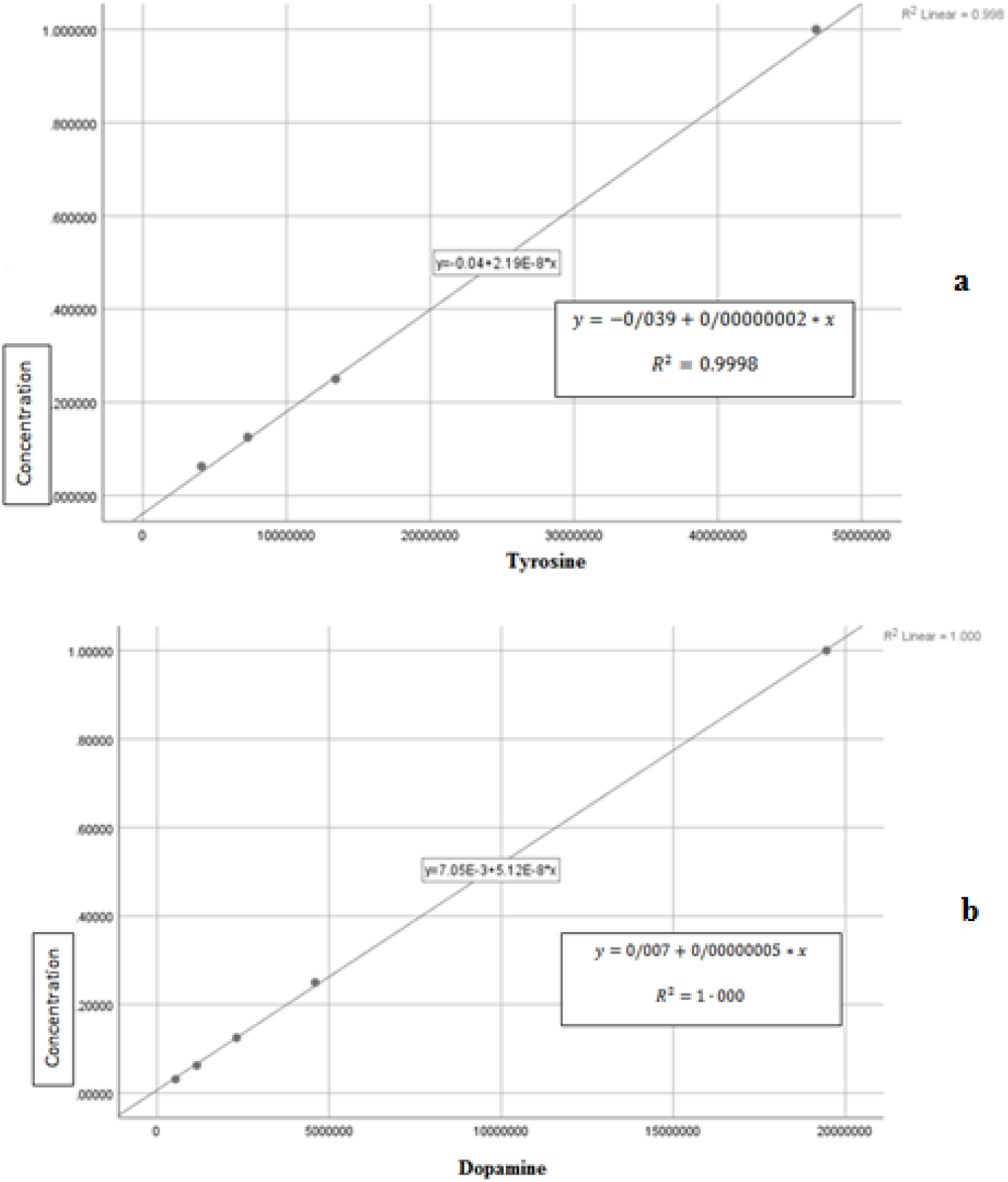
Calibration curves: a) tyrosine, b) dopamine

**Figure 4.**
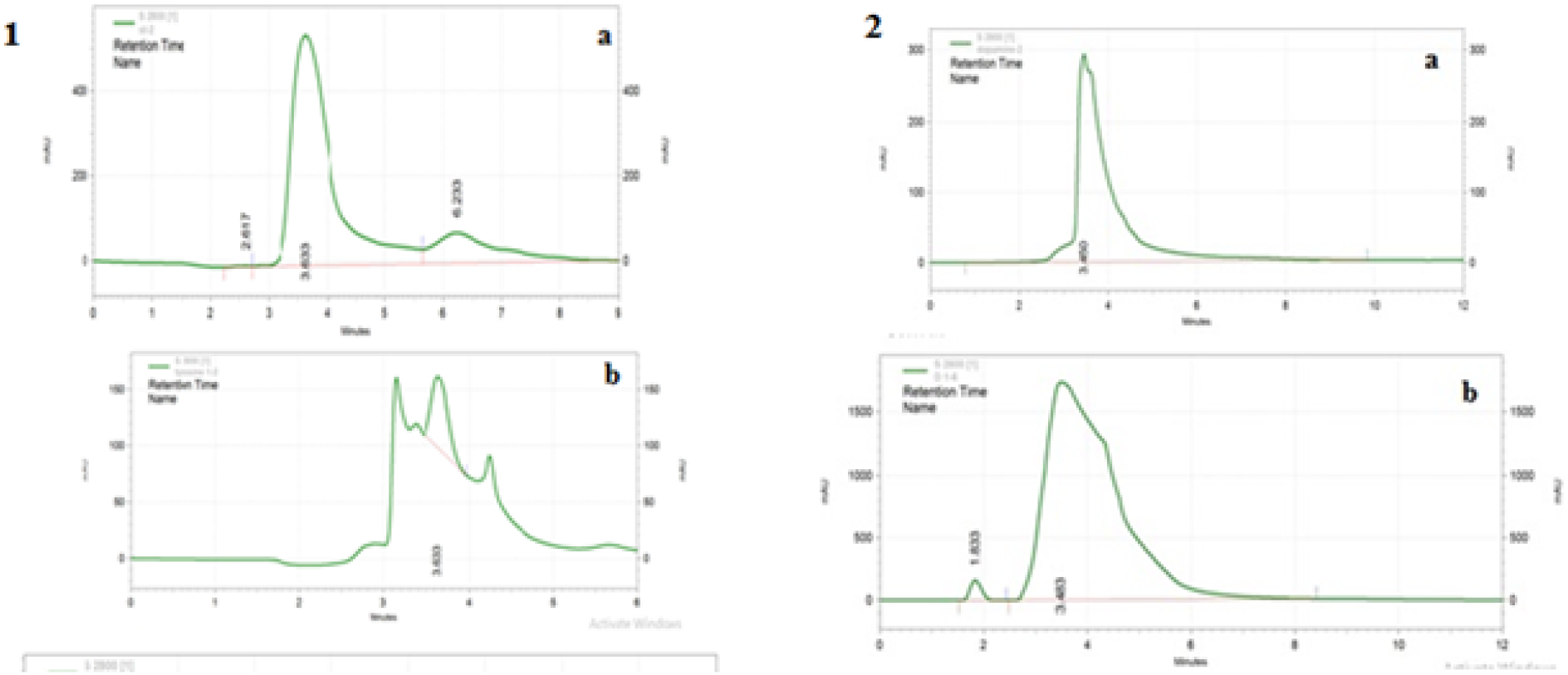
1. Tyrosine curves: a) standard (0.5 μgr/ ml), b) tyrosine serum level (at days 40) 2. Dopamine curves: a) standard (0.5 μgr/ ml), b) dopamine serum level (at days 40)

### Tyrosine serum level at different days after parasite injection

The obtained results regarding tyrosine showed that the tyrosine serum level is only measurable at 40 days (Table 1). At days 50, 60, 70, and 80 the tyrosine serum level was not detectable (Table 1 and Diagram 1).

**Table 1.** Tyrosine serum level

| Days after parasite injection | Sample 1 | Sample 2 | Sample 3 | Sample 4 | Sample 5 | Sample 6 | Sample 7 | Sample 8 | Tyrosine positive controls |
| --- | --- | --- | --- | --- | --- | --- | --- | --- | --- |
| 40 | 0.0532 | 0.0000 | 0.0000 | 0.0427 | 0.0000 | 0.0000 | 0.0000 | 0.0000 | 0.3003 |
| 50 | 0.0000 | 0.0000 | 0.0000 | 0.0000 | 0.0000 | 0.0000 | 0.0000 | 0.0000 | 0.3020 |
| 60 | 0.0000 | 0.0000 | 0.0000 | 0.0000 | 0.0000 | 0.0000 | 0.0000 | 0.0000 | 0.2121 |

**Diagram 1.**
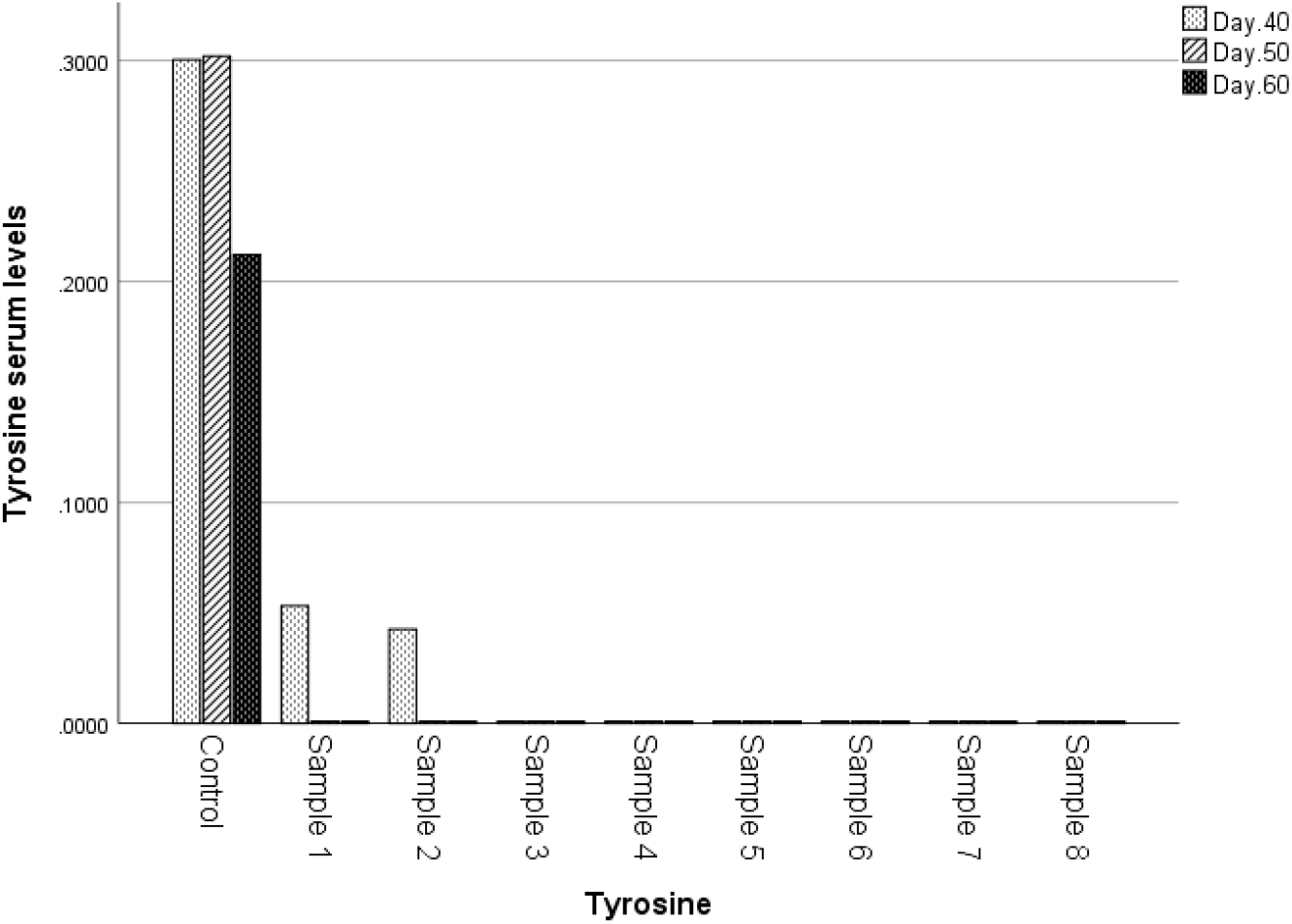
Descriptive statistics analysis of tyrosine level

**Diagram 2.**
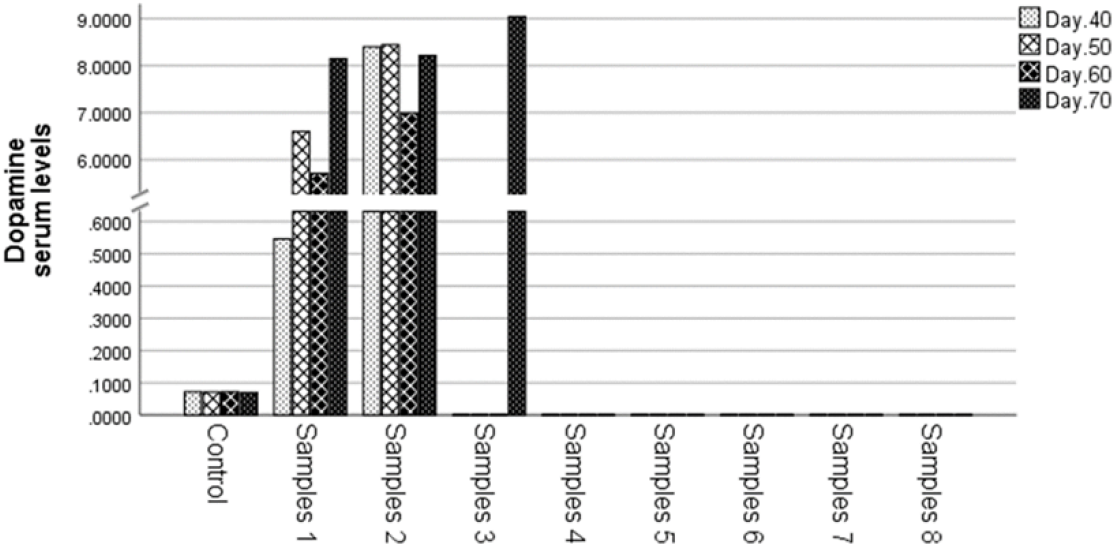
Descriptive statistics analysis of dopamine level

### Dopamine serum level at different days after parasite injection

The obtained results regarding dopamine showed that the dopamine serum level at days 40, 50, 60, and 70 had an approximately increasing trend after day 40. Dopamine serum level was not detectable at day 80 (Table 2).

**Table 2.** Dopamine serum level

| Days after parasite injection | Sample 1 | Sample 2 | Sample 3 | Sample 4 | Sample 5 | Sample 6 | Sample 7 | Sample 8 | Dopamine positive controls |
| --- | --- | --- | --- | --- | --- | --- | --- | --- | --- |
| 40 | 0.5459 | 0.0000 | 0.0000 | 8.4029 | 0.0000 | 0.0000 | 0.0000 | 0.0000 | 0.0716 |
| 50 | 6.5960 | 0.0000 | 8.4446 | 0.0000 | 0.0000 | 0.0000 | 0.0000 | 0.0000 | 0.0708 |
| 60 | 0.0000 | 5.7102 | 0.0000 | 0.0000 | 0.0000 | 6.9908 | 0.0000 | 0.0000 | 0.0718 |
| 70 | 0.0000 | 9.0439 | 0.0000 | 8.2153 | 0.0000 | 0.0000 | 0.0000 | 0.0000 | 0.0700 |

## Discussion

The results of the study showed that the level of dopamine had an approximately increasing trend after day 40. The results of a previous study revealed that the concentration of dopamine does not increase in infected animal models with acute toxoplasmosis (unpublished); however, the level of this hormone can be increased up to 14% in infected animal models with chronic toxoplasmosis. Although the cerebral level of dopamine was measured, reporting an increasing trend within the time regarding dopamine serum level in our results can be to some extent, in line with the results of this study that was done by Stibbs *et al.* (21). Another study in recent years has shown a high level of dopamine pertains to the presence of toxoplasmosis (22).

Based on our obtained results, low concentration of tyrosine at days 40 and increase of dopamine in most of the sample groups may confirm the previously obtained results regarding expression of encoding genes of tyrosine hydroxylase in *Toxoplasma* parasite. The authors of a study in 2009 believed that tyrosine level in infected animal models with toxoplasmosis decreased at two days after parasite injection due to the presence and expression of the encoding genes of tyrosine hydroxylase (AaaH1 and AaaH2) in *Toxoplasma* parasite (16). However, Alfonso *et al.,* silenced the AaaH2 gene and showed that other mechanisms probably are involved in decrease of tyrosine and increase of dopamine level in infected animal models with toxoplasmosis (23). So, the production of dopamine from tyrosine due to the tyrosine hydroxylase enzyme remains controversial.

Since in our study we could not detect the tyrosine level after day 40, we could not interpret our tyrosine data in comparison to the reported results in previous studies. No detection of tyrosine level in most of the sample groups probably referred to the very low concentration of tyrosine level in sera or low sensitivity of the HPLC system.

As mentioned, the relationship of toxoplasmosis and neurodegenerative disorders including Parkinson, Multiple Sclerosis and Alzheimer has been a discussable issue in recent years (7-10). The presence of *Toxoplasma* parasite in CNS and especially in the host’s brain can affect the transduction of neurotransmitter signals. This concept can be considered as the main reason for the behavioral changes in infected host with toxoplasmosis. Alteration of the tyrosine and dopamine serum and cerebral level can pertain to the varied behavioral disorders. In previous investigations, few studies attempted to improve the perception of the relation between neurotransmitter hormones and toxoplasmosis (16, 24).

To provide insights related to the alteration of the brain neurotransmitters level in animal models with toxoplasmosis, firstly we established a chronic toxoplasmosis model using PRU-II strain in BALB/c mice. *T. gondii* type I strains induce acute toxoplasmosis with different aspects ranging from aborted to severe forms and finally leads to the death of mice. Thus, to keep the mice alive and cyst formation in tissues, following the results of this study we used the PRU-II strain (25). Detection of antibodies in the sera of pilot mice using MAT, and also confirmation of cyst formation using microscopic and molecular methods, showed the ability of *T. gondii* PRU-II strain in inducing chronic toxoplasmosis in *in vivo* as well.

It seems that the results of this study can more confirm the changes that can happen in neurotransmitter signals in infected hosts with toxoplasmosis. This mechanism can be considered in the treatment of neurodegenerative disorders in future.

## Conclusion

This primary study may confirm the results of the previous studies regarding the relationship of toxoplasmosis and neurodegenerative disorders and also open a new venue for further investigations in this regard. It seems the evaluation of tyrosine and dopamine serum level using more samples and varied techniques might improve our obtained data in future studies. Also, the tyrosine and dopamine serum level can be compared with cerebral tyrosine and dopamine in infected animal models with chronic toxoplasmosis.

## Acknowledgment

This work was supported by Shiraz University of Medical Sciences, grant number [16519].

## References

1- Carruthers VB, Suzuki Y. Effects of Toxoplasma gondii infection on the brain. Schizophrenia bulletin. 2007; 33(3): 745–51.

2- Jones JL, Lopez A, Wilson M, Schulkin J, Gibbs R. Congenital toxoplasmosis: a review. Obstetrical & gynecological survey. 2001; 56(5): 296–305.

3- Saadatnia G, Golkar M. A review on human toxoplasmosis. Scandinavian journal of infectious diseases. 2012; 44(11): 805–14.

4- Dubey J, Tiao N, Gebreyes W, Jones J. A review of toxoplasmosis in humans and animals in Ethiopia. Epidemiology & Infection. 2012; 140(11): 1935–8.

5- Akstein RB, Wilson LA, Teutsch SM. Acquired toxoplasmosis. Ophthalmology. 1982; 89(12): 1299–302.

6- Ho Y-C, Sun H-Y, Chen M-Y, Hsieh S-M, Sheng W-H, Chang S-C. Clinical presentation and outcome of toxoplasmic encephalitis in patients with human immunodeficiency virus type 1 infection. J Microbiol Immunol Infect. 2008; 41(5): 386–92.

7- Lim SL, Rodriguez-Ortiz CJ, Kitazawa M. Infection, systemic inflammation, and Alzheimer’s disease. Microbes and infection. 2015; 17(8): 549–56.

8- Parlog A, Schlüter D, Dunay IR. Toxoplasma gondii-induced neuronal alterations. Parasite immunology. 2015; 37(3): 159–70.

9- Miman O, Kusbeci OY, Aktepe OC, Cetinkaya Z. The probable relation between Toxoplasma gondii and Parkinson’s disease. Neuroscience letters. 2010; 475(3): 129–31.

10- Oruc S, Karakaya F, Demirbas H, Cecen I, Küsbeci ÖY, Yaman M, et al. Relationship of Toxoplasma Gondii Exposure with Multiple Sclerosis. European Journal of General Medicine. 2016; 13(1).

11- Lagrue C, Poulin R. Manipulative parasites in the world of veterinary science: implications for epidemiology and pathology. The Veterinary Journal. 2010; 184(1): 9–13.

12- Lefevre T, Adamo SA, Biron DG, Misse D, Hughes D, Thomas F. Invasion of the body snatchers: the diversity and evolution of manipulative strategies in host–parasite interactions. Advances in parasitology. 2009; 68: 45–83.

13- Berdoy M, Webster JP, Macdon ald DW. Fatal attraction in rats infected with Toxoplasma gondii. Proceedings of the Royal Society of London Series B: Biological Sciences. 2000; 267(1452): 1591–4.

14- Frenkel J, Nelson B, Arias-Stella J. Immunosuppression and toxoplasmic encephalitis: clinical and experimental aspects. Human pathology. 1975; 6(1): 97–111.

15- Voller A, Bidwell D, Bartlett A, Fleck D, Perkins M, Oladehin B. A microplate enzyme-immunoassay for toxoplasma antibody. Journal of clinical pathology. 1976; 29(2): 150–3.

16- Gaskell EA, Smith JE, Pinney JW, Westhead DR, McConkey GA. A unique dual activity amino acid hydroxylase in Toxoplasma gondii. PloS one. 2009; 4(3): e4801.

17- Prandovszky E, Gaskell E, Martin H, Dubey J, Webster JP, McConkey GA. The neurotropic parasite Toxoplasma gondii increases dopamine metabolism. PloS one. 2011; 6(9): e23866.

18- Shaapan R, El-Nawawi F, Tawfik M. Sensitivity and specificity of various serological tests for the detection of Toxoplasma gondii infection in naturally infected sheep. Veterinary Parasitology. 2008; 153(3-4): 359–62.

19- Nakanuma Y, Ohta G. Histometric and serial section observations of the intrahepatic bile ducts in primary biliary cirrhosis. Gastroenterology. 1979; 76(6): 1326–32.

20- Asgari Q, Mohammadpour I, Pirzad R, Kalantari M, Motazedian MH, Naderi S. Molecular and Serological Detection of Toxoplasma gondii in Stray Cats in Shiraz, South-central, Iran. Iranian Journal of Parasitology. 2018; 13(3): 430.

21- Stibbs H. Changes in brain concentrations of catecholamines and indoleamines in Toxoplasma gondii infected mice. Annals of Tropical Medicine & Parasitology. 1985; 79(2): 153–7.

22- Flegr J. How and why Toxoplasma makes us crazy. Trends in parasitology. 2013; 29(4): 156–63.

23- Afonso C, Paixão VB, Klaus A, Lunghi M, Piro F, Emiliani C, et al. Toxoplasma-induced changes in host risk behaviour are independent of parasite-derived AaaH2 tyrosine hydroxylase. Scientific reports. 2017; 7(1): 13822.

24- Webster JP, McConkey GA. Toxoplasma gondii-altered host behaviour: clues as to mechanism of action. Folia parasitologica. 2010; 57(2): 95.

25- Xiao J, Li Y, Prandovszky E, Kannan G, Viscidi RP, Pletnikov MV, et al. Behavioral abnormalities in a mouse model of chronic toxoplasmosis are associated with MAG1 antibody levels and cyst burden. PLoS neglected tropical diseases. 2016; 10(4): e0004674.

